# X-ray phase contrast imaging of *Vitis* spp. buds shows freezing pattern and correlation between volume and cold hardiness

**DOI:** 10.1101/647248

**Authors:** Alisson P. Kovaleski, Jason P. Londo, Kenneth D. Finkelstein

## Abstract

Grapevine (*Vitis* spp.) buds must survive winter temperatures in order to resume growth when suitable conditions return in spring. They do so by developing cold hardiness through deep supercooling, but the mechanistic process of supercooling in buds remains largely unknown. Here we use synchrotron X-ray phase contrast imaging to study cold hardiness-related characteristics of *V. amurensis, V. riparia*, and *V. vinifera* buds: time-resolved 2D imaging was used to visualize freezing; and microtomography was used to evaluate morphological changes during deacclimation. Bud cold hardiness was determined (low temperature exotherms; LTEs) using needle thermocouples during 2D imaging as buds were cooled with a N_2_ gas cryostream. Resolution in 2D imaging did not allow for ice crystal identification, but freezing was assessed due to movement of tissues coinciding with LTE values. Freezing was observed to propagate from the center of the bud toward the outer bud scales. The freezing events observed lasted several minutes. Additionally, loss of supercooling ability appears to be correlated with increases in bud tissue volume during the process of deacclimation, but major increases in volume occur after most of the supercooling ability is lost, suggesting growth resumption processes are limited by deacclimation state.

**Highlight:** X-ray phase contrast imaging shows freezing occurs over several minutes and propagates from center toward tip of *Vitis* spp. buds. Incremental increase in bud volume correlates with cold deacclimation

## Introduction

Grapevines (*Vitis* spp.) produce compound mixed buds that contain both vegetative and reproductive tissue in a primary bud, and vegetative tissue in secondary and tertiary buds (Pratt, 1971). These buds are produced during the growing season, and transition into a dormant state to survive unsuitable growth conditions, such as drought or low temperature. Throughout the winter, grapevine buds will remain dormant and develop cold hardiness in order to prevent damage from low temperatures. Winter dormancy status is transitional, subtly changing from an endodormant to ecodormant status. During endodormancy, buds are recalcitrant to growth due to unknown internal repression. However, upon progressive chill accumulation, buds become ecodormant and will resume growth if exposed to permissive conditions (Lang *et al.*, 1987).

Growth resumption under forcing conditions, marked by the appearance of budbreak, is typically used to evaluate the changes in dormancy level that occur during winter (e.g., Londo and Johnson, 2014; Londo and Kovaleski, 2019). However, this comparison of phenological stage is dependent on comparable development between genotypes or species: if growth and expansion in the bud during dormancy release is not the same in all genotypes, we could incorrectly describe the relationship of cold hardiness and budbreak phenology. For example, phenological scales for budbreak in grapevine are based on observations of *V. vinifera* buds (Coombe and Iland, 2005; Andreini *et al.*, 2009) and may incorrectly describe changes that occur in wild species. Recently, this dormancy transition has been shown to be observed through the gradual increase in rate of cold hardiness loss that occurs with chill accumulation over the winter season (Kovaleski et al., 2018). However, the relationship between the kinetics of the deacclimation process and budbreak is different for species within *Vitis*: buds of cultivated grapevine (*V. vinifera*) only begin showing a budbreak phenotype after the majority of the supercooling ability is lost, whereas buds of *V. riparia* may present budbreak prior to fully losing their cold hardiness (Kovaleski *et al.*, 2018). Therefore, exploring differences in morphological aspects of *Vitis* species buds during deacclimation may elucidate important differences in time to budbreak and its relation to deacclimation kinetics.

While the majority of methods for studying bud morphology is destructive (e.g., sectioning), non-destructive methods may give better insight into how the developmental process of early budbreak affects the perception of phenological progression. Tomography imaging is an underused technique in plant sciences (Staedler *et al.*, 2013) that may be an alternative for the study of morphological differences between buds. This technique has recently become a consolidated method for the study of xylem and wood characteristics (Mayo *et al.*, 2010; Sedighi Gilani *et al.*, 2014; Cochard *et al.*, 2015; Torres-Ruiz *et al.*, 2015; Choat *et al.*, 2016; Malek *et al.*, 2016; Nardini *et al.*, 2017; Scoffoni *et al.*, 2017; Koddenberg and Militz, 2018), but other plant structures have also been imaged, such as developing maize seeds (Rousseau *et al.*, 2015), tomato leaves (Verboven *et al.*, 2015), and flowers and floral buds (Staedler *et al.*, 2013; Tracy *et al.*, 2017).

In higher latitudes, the dormant buds of woody species experience temperatures below the freezing point of water during the winter, and as a consequence have developed strategies to prevent bud death. Grapevine buds, as well as other woody perennials, survive these low temperatures and gain cold hardiness by promoting the supercooling of water in tissues (Burke *et al.*, 1976; Andrews *et al.*, 1984; Quamme, 1995). Through this process, pure water can remain liquid to temperatures close to −42 °C (Bigg, 1953). If the bud cold hardiness threshold is surpassed by low temperatures, bud mortality ensues, impairing growth and flowering in the following season. Maximum cold hardiness limits are different for different species, ranging from high negative temperatures (i.e., −7 °C) to very close to the ∼−42 °C supercooling limit (Quamme, 1995). Within grapevines, bud cold hardiness changes throughout the winter, primarily driven by changes in air temperature (Ferguson *et al.*, 2011, 2014; Londo and Kovaleski, 2017; Kovaleski *et al.*, 2018) and there is variation among species and cultivars within a species. Maximum cold hardiness has been observed to be mostly between −24 °C and −35 °C (Andrews *et al.*, 1984; Ferguson *et al.*, 2011, 2014; Londo and Kovaleski, 2017), with cultivated varieties being less cold hardy than wild species. Although low temperatures are the most limiting factor in plant distribution (Parker, 1963), the process through which plants control the supercooling point of buds and other structures remains largely unknown.

Damage in grapevine buds when supercooling fails is hypothesized to occur from the formation of intracellular ice (Andrews *et al.*, 1984; Mills *et al.*, 2006), however location of ice nucleation has not been studied in grapevines. The observation of the freezing process is the best means for understanding how the event causes damage (Molisch, 1897) and the identification of regions of the bud where supercooling fails can help understanding how plants control supercooling. Multiple techniques have been used to observe or infer ice formation in food and biological samples: indirect observation through freeze-substitution, identifying holes left in tissues by ice (Bevilacqua *et al.*, 1979); light microscopy (Molisch, 1897; Morris *et al.*, 1986; Guenther *et al.*, 2006; Stott and Karlsson, 2009; Endoh *et al.*, 2009, 2014); fluorescence microscopy with the aid of a microslicer for 3D ice structure (Do et al., 2004); NMR microscopy (Ishikawa *et al.*, 1997; Kerr *et al.*, 1998); freeze fracture Cryo-SEM (Endoh *et al.*, 2009, 2014); infrared imaging (Wisniewski *et al.*, 1997; Workmaster and Palta, 1999; Bauerecker *et al.*, 2008; Livingston *et al.*, 2018; Neuner *et al.*, 2019); and confocal laser scanning microscopy (Baier-Schenk *et al.*, 2005). In plants, Endoh *et al.* (2009) used light microcopy to examine extracellular ice crystals and freeze fracture Cryo-SEM to evaluate the presence of intracellular ice, based on the presence of crystalline ice vs. amorphous ice inside cells in buds of larch (*Larix kaempferi*). These methods, however, do not allow for temporal imaging of ice fronts. Using time-resolved X-ray phase contrast imaging, Sinclair *et al.* (2009) observed the growth of ice crystals in insect larvae in real time. X-ray phase contrast imaging of freezing appears to be an interesting option for imaging freezing in buds, considering the opaque nature of the structure, as well as the fact that it allows for temporal imaging of ice spreading (Sinclair *et al.*, 2009).

Understanding morphological changes within buds during deacclimation, as well as where freezing occurs may provide new insights into dormancy release and plant control over supercooling ability. Therefore, the objective of this study was to evaluate morphological development of buds from different *Vitis* species during loss of hardiness and budbreak, and image the freezing of buds to identify regions of the bud where the supercooling mechanism fails using X-ray phase contrast imaging.

## Materials and Methods

### Plant material and cold hardiness

Buds of *V. amurensis* PI588641, *V. riparia* PI588711, and *V. vinifera* ‘Riesling’ were collected from the field on 31 January 2018, prepared into single node cuttings and placed in a 4 °C cold room in cups of water. In preparation for imaging, sets of buds were removed from the cold room and placed under forcing conditions (22 °C, 16h/8h light/dark) periodically to deacclimate. Buds were removed on 31 Jan, 2 Feb, 5 Feb, 7 Feb, and 11 Feb 2018 for *V. riparia* and *V. vinifera*; and 8 Feb and 11 Feb 2018 for *V. amurensis*. On 13 Feb 2018, cold hardiness of buds was determined and buds were moved back into cold room (4 °C), where they were maintained throughout the imaging period to minimize changes in cold hardiness and developmental stage (Kovaleski *et al.*, 2018). This sampling scheme provided us with buds at 0, 2, 6, 8, 11, and 13 days of deacclimation for *V. riparia* and *V. vinifera*, and 0, 2, 5 days for *V. amurensis*.

Cold hardiness was determined through differential thermal analysis (DTA), as represented by individual low temperature exotherm (LTE) of buds (Mills *et al.*, 2006). In summary, buds are excised from cane and placed on thermoelectric modules (TEM) in plates, which are then placed in a programmable freezer. The freezer is cooled at −4 °C hour^−1^, and changes in voltage due to release of heat by freezing of water is measured by the TEMs and recorded via Keithley data logger (Tektronix, Beaverton, OR) attached to a computer. Deacclimation rates were estimated using linear regression (Kovaleski *et al.*, 2018) using R (ver. 3.3.0, R Foundation for Statistical Computing). R was also used to produce all plots.

### X-ray phase contrast imaging

Buds attached to a piece of cane were held on a custom-made cylindrical holder with mounting putty. The holder was attached to a small goniometer mounted on a Huber 4-circle diffractometer. Imaging was performed in the C-line at the Cornell High Energy Synchrotron Source (CHESS, Cornell University, Ithaca, USA). The monochromatic beam was expanded to 7 mm × 7 mm at X-ray energy 15 KeV. The sample-detector distance used was optimized to 0.5 m. Phase-contrast is produced when majority unperturbed beam interferes with angular deviations in the wavefront caused by density variations in the sample (Socha *et al.*, 2007). X-rays were converted into visible light using a rare-earth doped GGG (Gd_3_Ga_5_O_12_) crystal plate and imaged using an Andor Neo CMOS camera with a 5x objective lens.

For morphological changes in buds, tomographic-like imaging was performed with camera resolution of approximately 5 μm, obtained by adapting the objective lens. Buds were scanned while rotating over 180°, with images collected every ¼ or ½ degree. Reconstruction of bud structure based on these datasets was performed using Octopus Reconstruction software (ver. 8.8.1, Inside Matters, Belgium). After reconstruction, buds were visualized in 3D using OsiriX imaging software (ver. 8.0.1, Pixmeo, Switzerland). For volume measurements, a threshold was visually established for each bud to remove noise. The bud cushion (i.e., undifferentiated tissue connecting bud to shoot) was removed from the image, and only the bud itself was used. Volume was determined by counting the number of voxels in the 3D image using the ROI tool within OsiriX. Volume was observed as percent increase in volume (ΔV) from the sample in day 0. If more than one bud was imaged for day 0, the average volume of samples was used as the base value.

Freezing of the buds was observed using 2D time-lapse imaging with images at 2 μm pixel size. A 1 second exposure was used, but image capturing time effectively resulted in 0.56 Hz frequency. During the imaging, buds were cooled using a N_2_ gas cryostream (Oxford Cryosystems, UK), with a cooling rate of ∼−40 °C hour^−1^. A thermocouple in a 33-gage needle probe (Omega Engineering, Inc., USA) was inserted in the bud during imaging and used to measure the temperature inside the bud, and temperature measurements were recorded using an RDXL4SD data logger (Omega Engineering, Inc., USA). LTEs for these samples were observed as temperature deviations from the linear rate of cooling. Contraction of the mounting putty due to cooling resulted in a slow downward drift of the bud, therefore image sets where aligned using the Linear Stack Alignment with SIFT plugin (https://imagej.net/Linear_Stack_Alignment_with_SIFT; Lowe, 2004) in Fiji (ImageJ ver. 2.0.0; Schindelin *et al*., 2012), and then cropped to remove black edges. Kymographs were obtained from the aligned image stacks using Fiji. To evaluate changes in buds over time, multi-scale structural similarity (MS SSIM) index (Wang *et al.*, 2004) was quantified between each image and the initial image using the MS SSIM index plugin (https://imagej.nih.gov/ij/plugins/mssim-index.html) in Fiji. To compare different regions of the buds, MS SSIM index values were normalized to a maximum of 100% and minimum of 0%. Image stacks were transformed into videos using Fiji, and temperature and image information were matched using time stamps in data-logger and images, inserted using the *Series Labeler* plugin (https://imagej.net/Series_Labeler). The time required to image buds for 3D scans and freezing scans limited the number of samples.

## Results

Deacclimation of the buds was well described by linear behavior until the limits of detection of LTEs (Fig. 1). *V. amurensis* and *V. riparia* had similar deacclimation rates, at 2.24 °C day^−1^ (R^2^= 0.89) and 2.12 °C day^−1^ (R^2^= 0.92), respectively. *V. vinifera* had a lower deacclimation rate, at 1.33 °C day^−1^ (R^2^= 0.95). LTEs determined using needle probes inserted in the buds during imaging of freezing produced similar results to those using the regular DTA method. In *V. riparia*, the last two time points were not used for rate determination as buds had already started to open.

**Figure 1.**
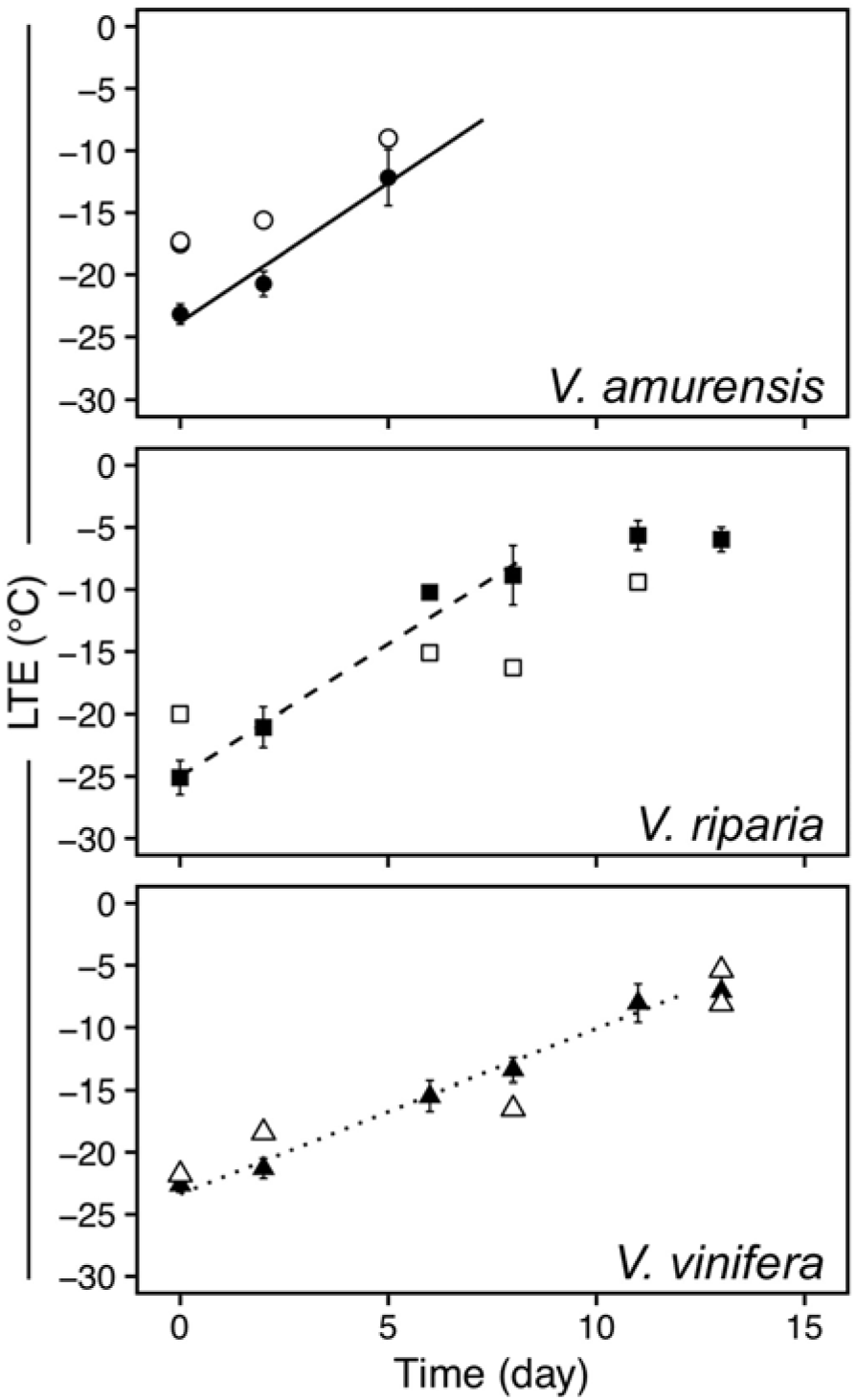
Deacclimation of *Vitis amurensis, V. riparia*, and *V. vinifera* ‘Riesling’ under forcing conditions. Full symbols represent average bud cold hardiness estimated through differential thermal analysis (DTA), while open symbols represent freezing temperature of single buds under a cryostream. Error bars represent standard deviation of the mean. Deacclimation rates (linear regression) were 2.24 °C day^−1^ (R^2^= 0.89), 2.12 °C day^−1^ (R^2^= 0.92), and 1.33 °C day^−1^ (R^2^= 0.95) for *V. amurensis, V. riparia*, and *V. vinifera*, respectively (*P*<0.001 for all), at 22 °C and 16h/8h light/dark.

Both the vegetative and reproductive aspects of the mixed Vitis bud structure were visible in micro-CT imaging (Figs. 2–4). As a consequence of faster deacclimation and development, *V. riparia* buds were imaged through a wider range of developmental stages (Fig. 2) than *V. vinifera* (Fig. 3). *V. riparia* was imaged in E-L stages 1 – “winter bud” (Fig. 2A–C), 2 – “bud scales opening” (Fig 2D), and 3 – “wooly bud” (Fig 2E). *V. vinifera* buds have an outer appearance of E-L stage 1 in Figs. 3A–C, and is at an early stage 2 in Figure 3D. With a reduced number of sampling dates, *V. amurensis* buds were all at E-L stage 1 and had the lowest range of development imaged (Fig. 4). Primary, secondary, and tertiary buds are visible in the still images shown for all three species. Images provide clear identification of inflorescences in the primary bud, even in the fully dormant state (day 0; Figs. 2A, 3A, 4A).

**Figure 2.**
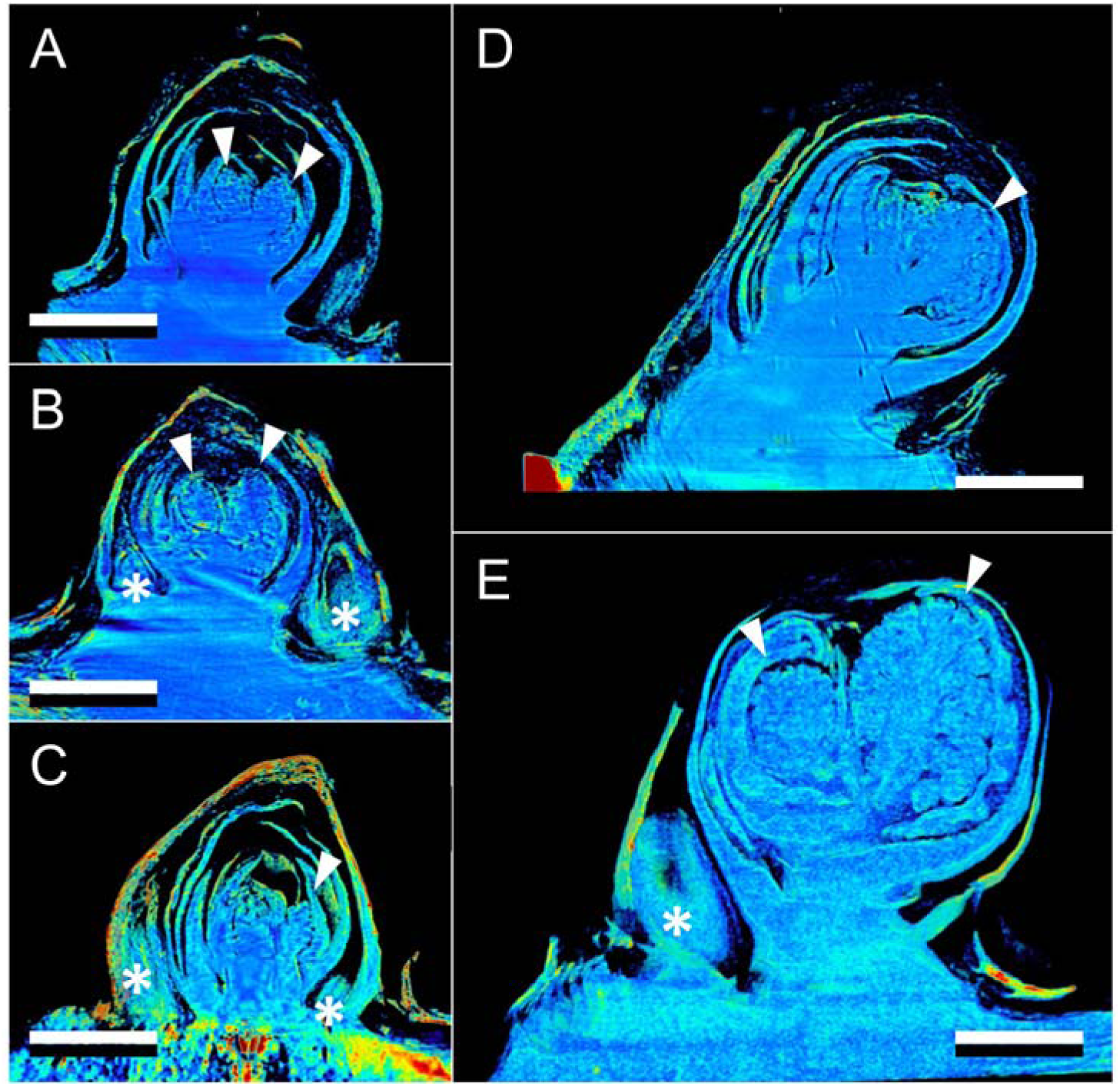
Development of *Vitis riparia* buds during budbreak reconstructed using X-ray microtomography. Buds shown were imaged at 0 (A), 2 (B), 8 (C), 11 (D), and 13 (E) days under forcing conditions. Full arrow heads indicate inflorescences, asterisks indicate secondary and tertiary bud. Scale bar = 1 mm (all images are in the same scale).

**Figure 3.**
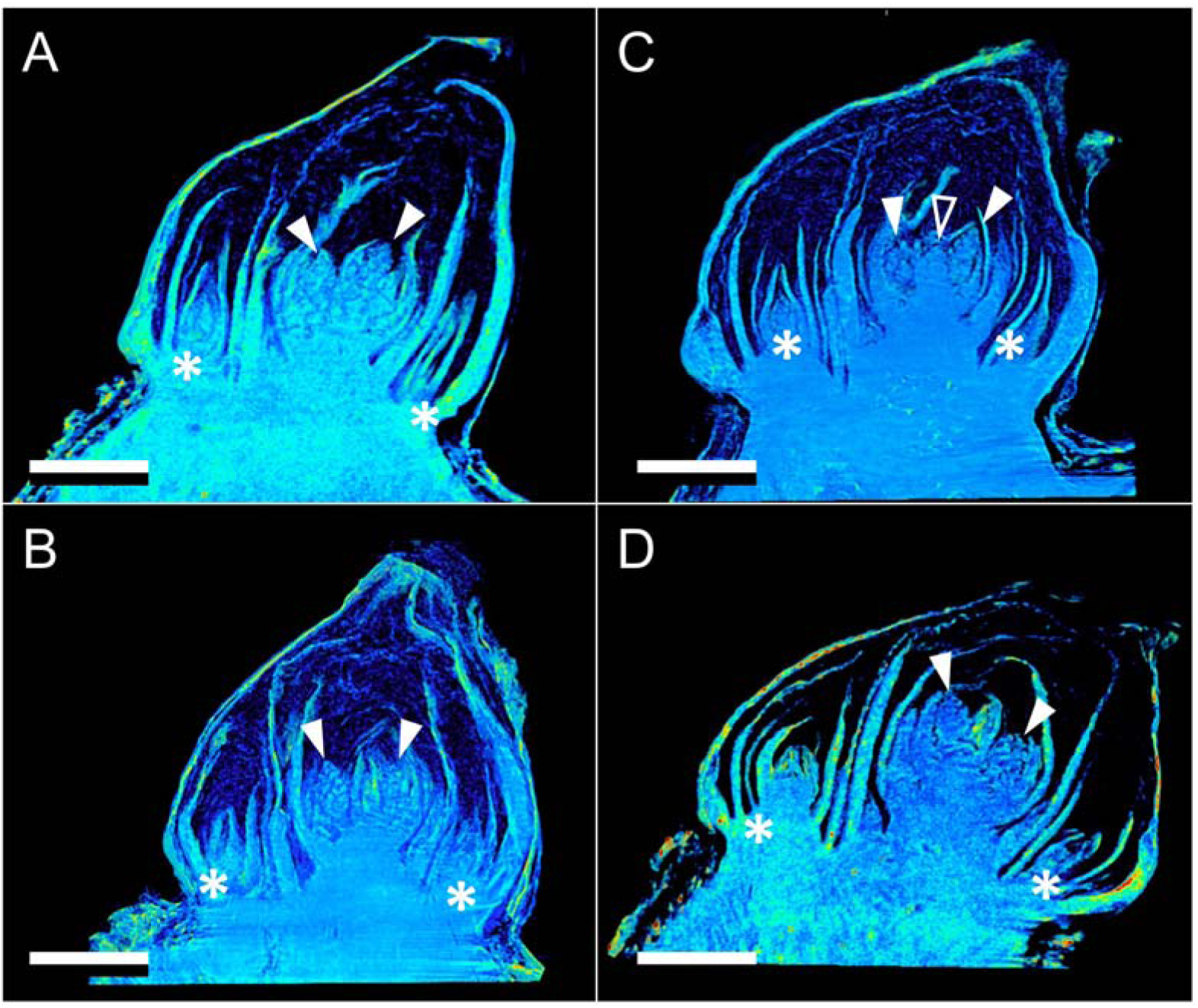
Development of *Vitis vinifera* buds during budbreak reconstructed using X-ray microtomography. Buds shown were imaged at 0 (A), 2 (B), 8 (C), and 13 (D) days under forcing conditions. Full arrow heads indicate inflorescences, open arrow head indicates apical meristem, asterisks indicate secondary and tertiary bud. Scale bar = 1 mm (all images are in the same scale).

**Figure 4.**
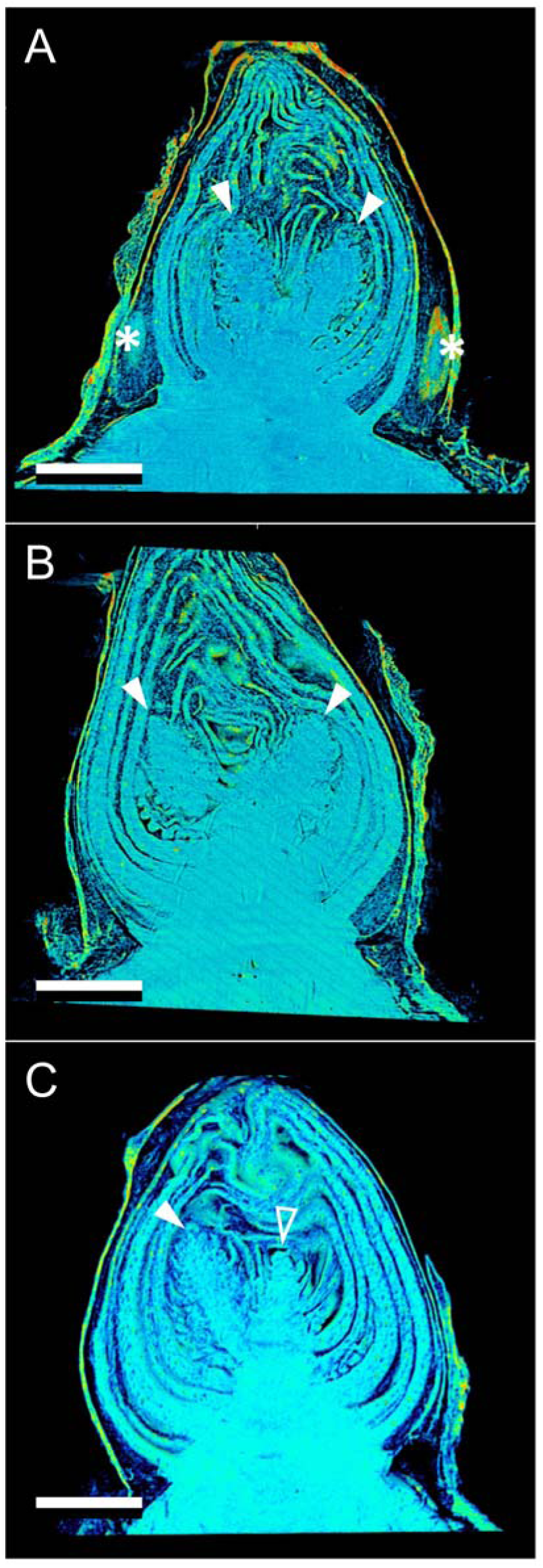
Development of *Vitis amurensis* buds during budbreak reconstructed using X-ray microtomography. Buds shown were imaged at 0 (A), 2 (B), and 5 (C) days under forcing conditions. Full arrow heads indicate inflorescences, open arrow head indicates apical meristem, asterisks indicate secondary and tertiary bud. Scale bar = 1 mm (all images are in the same scale).

Clear morphological differences can be seen when comparing buds of the different species. *V. riparia* buds are much smaller than *V. vinifera* and *V. amurensis*. The inflorescence primordia in *V. riparia*, however, take up much more of the volume of day 0 buds in *V. riparia* than in *V. vinifera*. Both *V. vinifera* and *V. riparia* have inflorescence primordia of ∼0.5 mm, whereas in *V. amurensis* they are ∼1mm long and appear more developed. Buds of *V. vinifera* have much more space between the leaf primordia, inflorescence primordia, and the outer bud scales compared to the two other species, especially *V. amurensis*. This space is occupied by “wool” or “hair”, most visible in Figs. 1B and C. *V. amurensis* buds are very compact at the dormant stage, and there is very little space between the scales and leaf primordia, which can be seen folding down on the top, as if constrained by the outer scales (Fig. 4A).

In *V. riparia*, there are very little differences in morphology between the buds until day 8 (Figs. 2A–C). However, once LTE values were above –10 °C (close to the limit of detection where HTEs and LTEs may combine in DTA; Figure 1), a noticeable increase in the bud size can be observed (Figs. 2D and E, 5). Much of this change appears to be due to the expansion and development of the inflorescence primordia, and elongation of the base of the primary bud (shoot). In *V. vinifera*, the inflorescence primordia appear to remain the same size as the buds lose hardiness but there is a noticeable expansion of the base of the primary bud. In *V. amurensis*, there are no clear internal differences seen between day 0 and 5.

The visual assessments of expansion in bud tissues are confirmed by analysis of the volume of tissue (ΔV) in the buds (Fig. 5). *V. riparia* buds reached the greatest expansion in volume within the time analyzed, reaching at day 13 almost triple the size of buds in day 0. *V. amurensis* appears to have a similar slope when the first days are considered compared to *V. riparia*, while *V. vinifera* has the slowest increase in bud volume. Both *V. riparia* and *V. vinifera* buds had increased ∼50% in volume when most of the hardiness was lost (day 8 and day 13, respectively), although the rate of volume increase is much higher after all cold hardiness is lost for *V. riparia* (day 8-13). Pearson’s correlation for LTE and ΔV for *V. amurensis, V. riparia* and *V. vinifera* are 0.94, 0.96 and 0.96, respectively.

**Figure 5.**
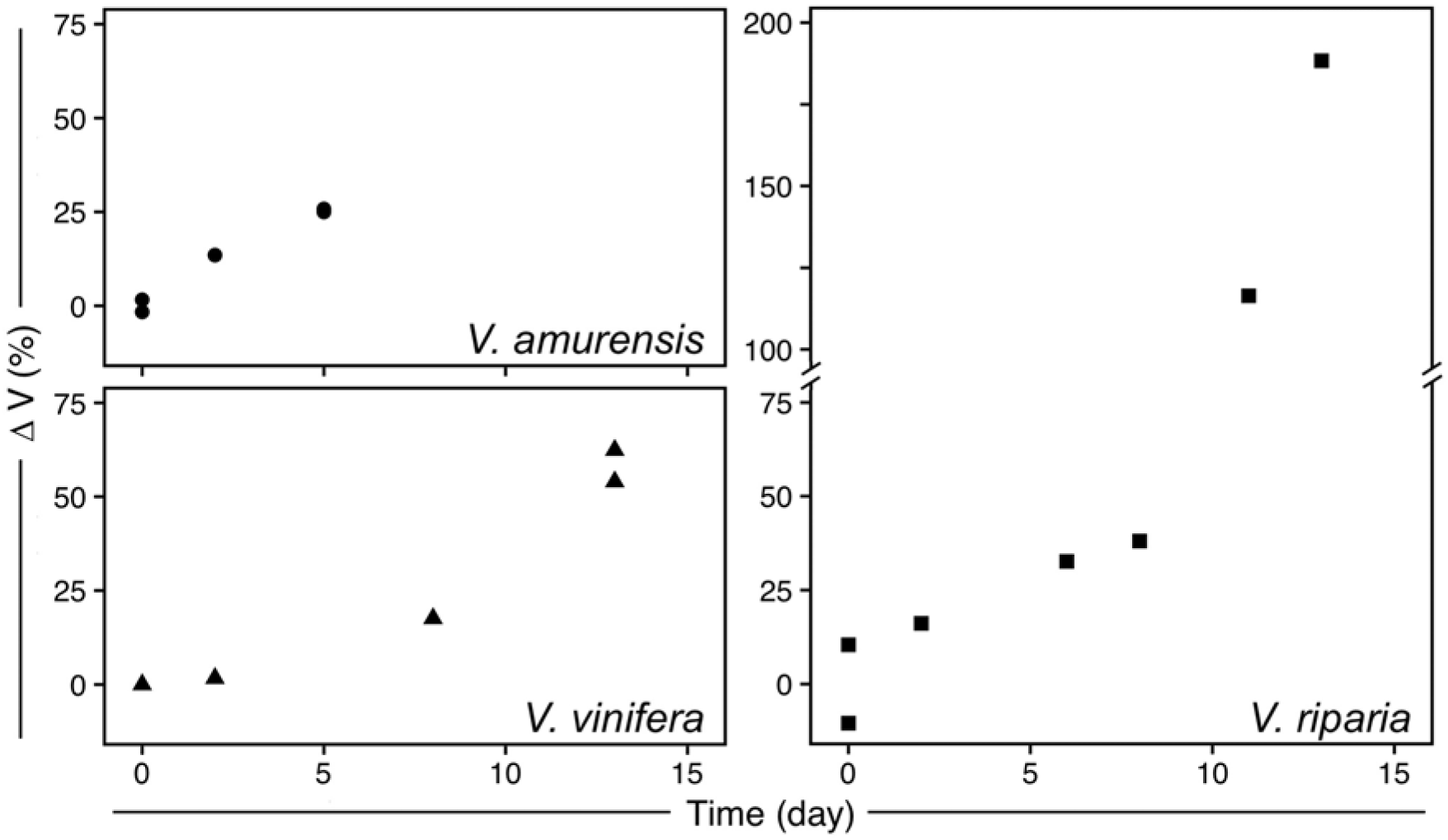
Increase in volume (ΔV) of *Vitis amurensis, V. riparia*, and *V. vinifera* during deacclimation. Volume was determined by counting the number of voxels in X-ray microtomography-reconstructed buds, therefore not including air space. ΔV wascalculated as the percent increase in volume from sample (or average of samples) at day 0.

Freezing of the buds occurred from the inside-out (Fig. 6, Supplementary Figure S1, Supplementary Videos S1–S3). The videos are produced based on projection images that show the accumulated structure of buds (Fig. 6A – same bud as in Fig. 2D). Freezing in this *V. riparia* bud is observed at −9.4 °C / 28:41 mm:ss (Supplementary Video S1). In a kymograph taken through the mid-section of the bud, an expansion of tissues is visualized by the drift outward in the structures (Fig. 6B). When aligning the MS SSIM and temperature probe data (Figs. 6C and D), we observe that there is a slow decay in MS SSIM during the initial cool down. The fastest decay in MS SSIM occurs simultaneously with the recording of increase in temperature due to heat release of water freezing. When comparing a top section with a bottom section of the bud, the MS SSIM index decays to a minimum value earlier than in the top (Fig. 6C inset). In a *V. amurensis* bud from day 0 (Fig. 4A), freezing of the primary bud occurred at −17.5 °C [Supplementary Figure S1; Supplementary Video S2 (time-stamp 32:10)]. The freezing resulted in an increase in the inner temperature of the bud of ∼8 °C, reaching −9.4 °C. A much smaller increase in temperature occurs at 39:14, caused by the freezing of secondary bud. Evaluation of MS SSIM shows similar results in *V. amurensis* (Supplementary Figure S1). Detection of freezing in *V. vinifera* was subtler, with a very slight expansion of the center portion of the bud [Supplementary Video S3 (−18.3 °C / 33:30)]. The movement of tissues that signals freezing can be observed occurring over minutes in all genotypes: between 28:41 and ∼34:00 in *V. riparia*, 32:10 to ∼41:00 in *V. amurensis*, and 33:30 to ∼36:00 in *V. vinifera*.

**Figure 6.**
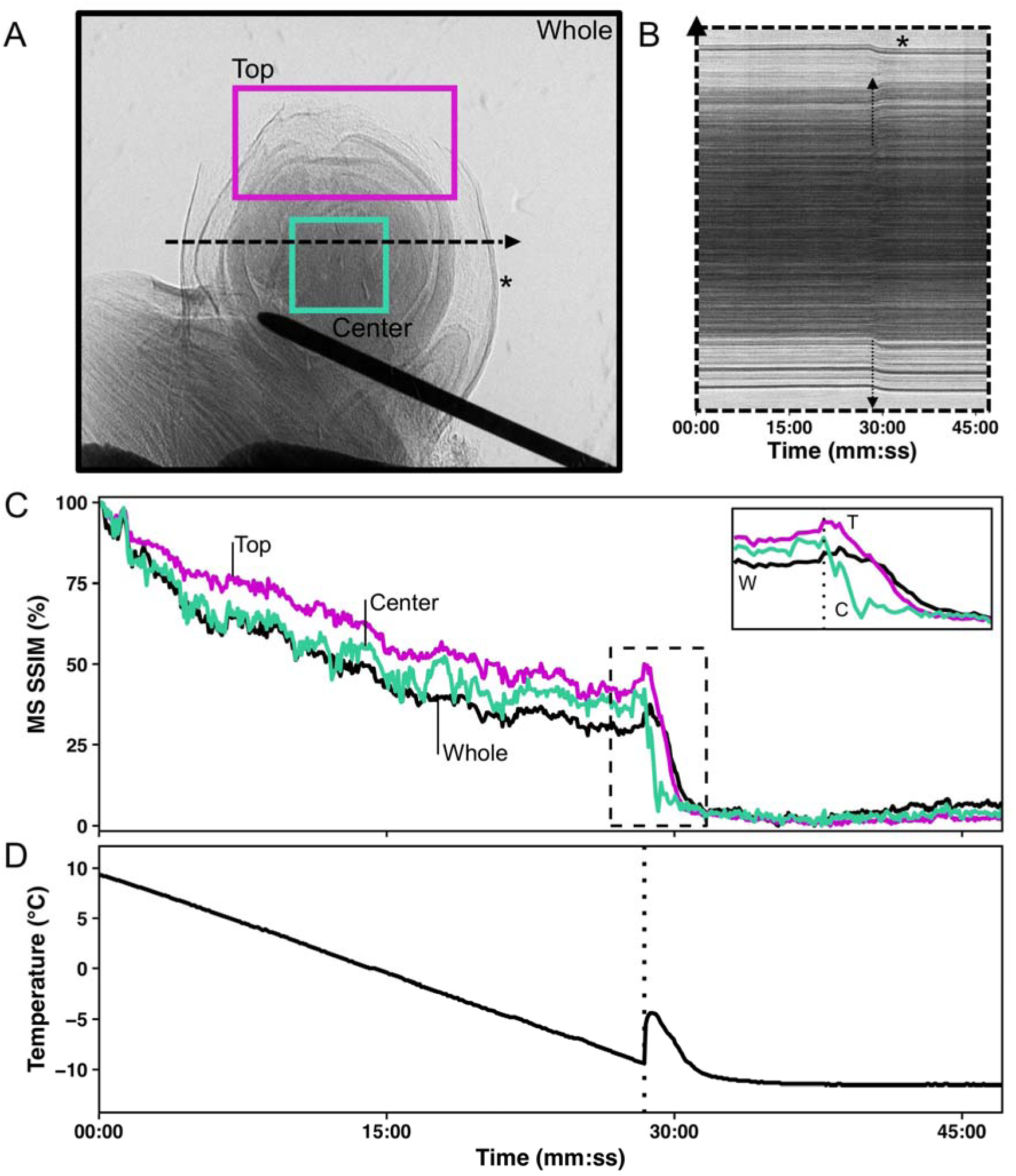
Characteristics of freezing in a bud of *Vitis riparia* after 11 days of deacclimation (see Supplementary Video S1). (A) Still image of bud at start of freezing; Black (whole image), magenta (top of the bud), and cyan (center of the bud) show areas analyzed; dashed line through center of the bud shows pixels used to build kymograph. (B) Kymograph resulting from line of pixels in the center of the bud; arrows show the start of freezing; asterisk marks the outer bud scale that moves inward. (C) Normalized multi-scale structural similarity index (MS SSIM) for three areas in (A); dashed box is shown expanded in the inset, dotted line marks the start of freezing event. (D) Temperature profile measured by thermocouple inside the bud; dotted line marks the start of freezing event.

## Discussion

Only recently has X-ray microtomography begun to be used for the exploration of floral development in annual plants (Tracy *et al*., 2017 and referencing papers), but here we demonstrate the use of this technique to study morphological changes in buds of woody perennials. More importantly, we used quantitative data derived from tomography scans to explore concepts related to cold hardiness, and X-ray phase contrast imaging to visualize freezing. We demonstrate that small gains in volume occur during deacclimation, but increases are much faster once most of the supercooling ability of buds is lost, suggesting that the supercooled state in some way limits growth and development in dormant grapevine buds. Although the freezing method and rate of cooling were different than that typically used, the use of temporal X-ray imaging clearly shows that the freezing of tissues occurs from the inside of the bud and propagates to the outside, and that the freezing of bud tissues can last several minutes.

The non-destructive nature of X-ray phase contrast imaging is an interesting aspect for study of supercooling in buds, where damage to structure can result in loss of the phenotype (Quamme *et al.*, 1995). Although long-term survivorship of the buds was not tested, and radiation levels could potentially lead to cell death (Socha *et al.*, 2007; Sinclair *et al.*, 2009), buds that were imaged showed LTEs in comparable levels to those determined in standard DTA analysis (open vs. full symbols in Fig. 1, respectively). This demonstrates that at least for a few hours (scan for 3D imaging was ∼1h, followed by the freezing scan) the buds remained viable, as dead buds show no, or much warmer, LTEs. This also demonstrates that the thermocouple probes in the needles are effective for the detection of exotherms related to cold hardiness of buds, and that placement of needles did not disrupt usual supercooling. The comparable LTE levels are very interesting considering two aspects: (i) the high rate of cooling used and (ii) the lack of observed high temperature exotherms (HTEs). The rate of cooling used in the cryostream was ∼10x higher than that normally used in DTA – including DTA measurements used here to determine initial cold hardiness. The higher rate was required due to the time constrains for beam access, and faster freezing allowed us to image a greater number of buds. While the rates of cooling at the level used in this study reportedly cause a decrease in LTE temperature (more negative) of *V. vinifera* hybrid grapevines (Quamme, 1986), higher rates of cooling result in freezing at warmer temperatures for *Rhododendrum* sp. (Ishikawa and Sakai, 1981). We did not observe a particular trend when all species are taken into account. However, all the buds of *V. amurensis* froze at higher temperatures than the expected. This is likely a result of the over 2x greater rate of deacclimation this species has compared to the other two at low temperatures (Kovaleski *et al.*, 2018), and therefore storage may have resulted in some cold hardiness loss.

Needle probe data did not show any HTEs in terms of temperature deviations from the linear rate of cooling. There were also no visible cues or MS SSIM deviations relative to HTEs in buds in phase contrast imaging (Fig. 6, Supplementary Figure S1, Supplementary Videos S1–S3). In regular DTAs, HTEs are enhanced by the use of water sprays (Mills *et al.*, 2006), resulting in much larger peaks than LTEs. It is possible, however, that the HTE signal in non-wetted buds comes from the piece of cane attached to the bud, rather than the extracellular space in the bud itself. This agrees with the report by Neuner *et al*. (2019) in the vast majority of the 37 species studied there was no ice within the buds even after HTE. HTEs may also be a result of condensation followed by freezing, or sublimation of water vapor on TEMs during the cooling in DTAs. The cryostream used in our setup has a ring of warmer, dry N_2_ gas surrounding the N_2_ cryostream, which prevents sublimation or condensation on the sample during the cooling. The possible HTE-corresponding signals seen were very slight lags in the temperature decrease (e.g., ∼−10.0 °C in Supplementary Figure S1D). This behavior might indicate extra-organ freezing happens in grapevines, without extracellular ice forming within primordia such as described in other species (Quamme *et al.*, 1995; Endoh *et al.*, 2009, 2014). If the HTE happens in tissues further from the center of the bud, it is possible that the placing of the needle inserted could prevent or diminish the perception of temperature changes caused by tissues away from the center of the bud. However, LTEs corresponding to secondary buds were seen and measured based on temperature changes (Supplementary Figure S1D). The lack of HTE may also be an artifact of the high rate of cooling used as compared to regular DTA. However, LTEs measured were not far from expected values, and therefore HTEs may not be necessary for supercooling to occur. Further testing using multiple needles in buds should be conducted using different methods and rates of cooling to verify the occurrence, location, and importance of HTEs.

The resolution obtained in the freezing images of 2 μm pixel size was not enough to resolve ice crystals in the buds such as observed by Sinclair *et al.* (2009) imaging larvae of *Chymomyza amoena* and *Drosophila melanogaster*. Larvae have free lymph in large volumes, allowing the formation of large crystals within their bodies. In grapevine buds, most of the water is located inside cells with diameter less than 20 μm. While imaging at a higher resolution (∼1 μm) is possible, increasing the resolution results in a smaller area of imaging (Verboven *et al.*, 2015) that would likely not fit a whole bud. Therefore, we assessed freezing as the movement resulting from volume expansion due to phase change in water, also observed by Sinclair *et al.* (2009).

Despite the cryostream hitting the bud from the top, possibly generating a small temperature gradient, freezing was directly observed to occur from the inside initially, followed by outward progression in all species (Fig. 6, Supplementary Figure S1, Supplementary Videos S1–S3). Based on the MS SSIM, we can see the decay occurs earlier in the center portion vs. the top portion (Fig. 6, Supplementary Figure S1). This is also very clearly observed in Supplementary Video S2, where the *V. amurensis* bud scales located in the distal portion of the bud appear to be the last ones to freeze as they slightly unfold. This is a similar behavior to what was described by Quamme *et al.* (1995) for buds of peach (*Prunus persica*), in which ice propagates from the subtending tissues into the bud. Considering the apparent higher cold hardiness in bud scales compared to the shoot tip area, future studies exploring cold hardiness may want to compare these structures within the bud in terms of anatomy and gene expression.

Clear morphological differences are seen between the three species studied. *V. vinifera* has much less green (solid) tissue per bud volume than the other two species analyzed. Much of the bud volume is actually occupied by wool material (most visible in Fig. 3B and C). This adaptation is potentially linked to the region of origin: buds of *V. vinifera* likely had to adapt to reduce water loss during the dormant season in a warmer and drier place (Mediterranean) as compared to the areas where *V. amurensis* and *V. riparia* are native to (Northeastern Asia and North America, respectively). The differences in tissue of *V. amurensis* and *V. riparia* buds compared to *V. vinifera* also validate visible differences observed during budbreak. *V. riparia* has faster early development in the E-L scale compared to *V. vinifera*, even when responses to temperature are corrected (Kovaleski *et al.*, 2018). This may be a result of the larger volume of green tissue present in buds of *V. riparia* compared to *V. vinifera*. The implication of this observation is that there is less bud volume available for expanding tissues to fill in *V. riparia*, thus bud scales are forced open “earlier” in this species. Although it was not seen, *V. amurensis* would probably have similar or earlier budbreak than *V. riparia*, considering all of the tissues within the bud are extremely compacted and any expansion might result in appearance of early stages of budbreak (opening of the outer scales). It is not clear however how these morphological differences may implicate in greater maximum cold hardiness in *V. amurensis* and *V. riparia* compared to *V. vinifera* (Londo and Kovaleski, 2017).

The increase in volume is positively correlated with deacclimation, and faster increase of volume and deacclimation rates are seen in *V. amurensis* and *V. riparia* as compared to *V. vinifera* (Fig 1 and 5). This could indicate that increases in volume are reducing the ability of buds to supercool, likely as a result of influx of water leading to turgor (Xie *et al.*, 2018). Although it is not known how plants are able to control levels of deep supercooling, from a physical aspect it is known that larger volumes of water are at higher risk of ice nucleation at any given temperature (Bigg, 1953). Cold hardiness is correlated with bud water relations (Ishikawa and Sakai, 1981; Richards and Bliss, 1986), and *V. vinifera* buds have an increase in ∼25% water content from dormant to budbreak stage (Xie *et al.*, 2018; Meitha *et al.*, 2018). However, it is important to acknowledge that metabolic changes within the bud during deacclimation can also play a part in the loss of supercooling ability (Meitha *et al.*, 2018). The more rapid increase in volume in the later stages may be a result of re-establishment of vascular connections between the bud and the cane (Xie *et al.*, 2018). Newly developed xylem does not appear clearly such as large vessels in the cane, but the use of contrasting agents (Staedler *et al.*, 2013) could be used to evaluate the formation of xylem connections such as is done with dyes and light microscopy (Xie *et al.*, 2018). Contrasting agents may also be of potential use to more easily segment different parts of the bud in a virtual histology approach if differential uptake by tissues leads to clear density differences (Rousseau *et al.*, 2015), which could be tested in future assessments.

Buds took several minutes to completely freeze. This occurred despite the steep cooling rate and the cooling method based on a cryostream, which would reduce the difference in air to bud temperature by greatly decreasing the boundary layer (Grace, 2006). This contradicts previous descriptions that the freezing that produces an LTE is sudden (Quamme, 1995), and lasts only a few seconds in buds of multiple species by Neuner *et al.* (2019) using infrared imaging for infrared DTA (IDTA). Because IDTA only observes the increase in temperature of the bud, propagating heat from the center of the bud to the outside would appear the same way as if ice was forming in those tissues. Indeed, our temperature probe data shows that the derivative of temperature measurements is only positive for a very brief period of time (Fig. 6D, Supplementary Figure S1D). However, both the MS SSIM (Fig. 6C, Supplementary Figure S1C) and Supplementary Videos demonstrate that the wave of bud freezing lasts longer, even as the downward trend in the temperature measurements has resumed. Such downward trend in temperature would not appear in the images from IDTA, and therefore a great portion of the time for freezing is ignored. It is also important to note that the freezing of the secondary bud in *V. amurensis* (Supplementary Figure S1; Supplementary Video S2) appears to be a separate event entirely. This suggests that the freezing of secondary buds is protected from the primaries by a barrier that is not overcome by the propagation of ice that occurs upon initial freezing.

There was a difference in the time it took to completely freeze different buds. The size difference and amount of green tissue between species and development stages might justify why some buds froze more quickly compared to the other species if a similar rate of intracellular ice growth propagation is considered (Acker *et al.*, 2001). Although *V. amurensis* has buds with more volume than *V. vinifera*, it is possible that the wool in *V. vinifera* buds, as well as the shape of it reduced the rate of heat loss to the exterior. Energy balance studies comparing theoretical buds may allow for explanations for the differences in the duration of freezing. However, it is unlikely that insulation capabilities of bud tissues would be an adaptive response to increase cold hardiness, since air temperature changes in nature occur at a much lower rate and low temperature exposure lasts for longer periods of time.

X-ray microtomography proved to be a useful approach to identify structures within a bud, as well as for quantitative analysis of changes during loss of cold hardiness and early budbreak. Although our setup required removal of the bud from the cane, adaptation of a sample holder could lead to observation of growth in the same bud during development. Future explorations with contrasting agents (Staedler *et al.*, 2013) may aid in anatomical studies, with special interest to water movement in the bud. High temperature exotherms were not visible or measurable, which indicates they may be an artifact of the larger sensors used in DTA. The use of 2D time-lapse X-ray phase contrast associated with a thermocouple was useful in identifying how ice spreads throughout the bud. We identified the differential response where the center of the bud is where ice nucleates and propagates from toward the scales, and showed that extra-organ freezing on scales or extracellular are not necessary for supercooling of buds of different grapevine species. Finally, ice propagation observed by movement of tissues occurred over several minutes.

## Supplementary data

**Fig. S1.** Characteristics of freezing in a bud of *Vitis amurensis* stored at 4 °C.

**Video S1.** X-ray phase contrast imaging of *Vitis riparia* bud during freezing.

**Video S2.** X-ray phase contrast imaging of *Vitis amurensis* bud during freezing.

**Video S3.** X-ray phase contrast imaging of *Vitis vinifera* bud during freezing.

### Acknowledgements

This work is based upon research conducted at the Cornell High Energy Synchrotron Source (CHESS) which is supported by the National Science Foundation under award DMR-1332208. This work was partially supported by: CAPES, Coordenação de Aperfeiçoamento de Pessoal de Nível Superior, Brazil, award number 12945/13-7; by the National Science Foundation Plant Genome Research Program Award 1546869; and by an appointment to the Agricultural Research Service (ARS) Research Participation Program administered by the Oak Ridge Institute for Science and Education (ORISE) through an interagency agreement between the U.S. Department of Energy (DOE) and the U.S. Department of Agriculture (ORISE is managed by ORAU under DOE contract number DE-SC0014664). All opinions expressed in this paper are the authors’ and do not necessarily reflect the policies and views of USDA, ARS, DOE, or ORAU|ORISE.

## Supplementary data

**Supplementary Figure S1.**
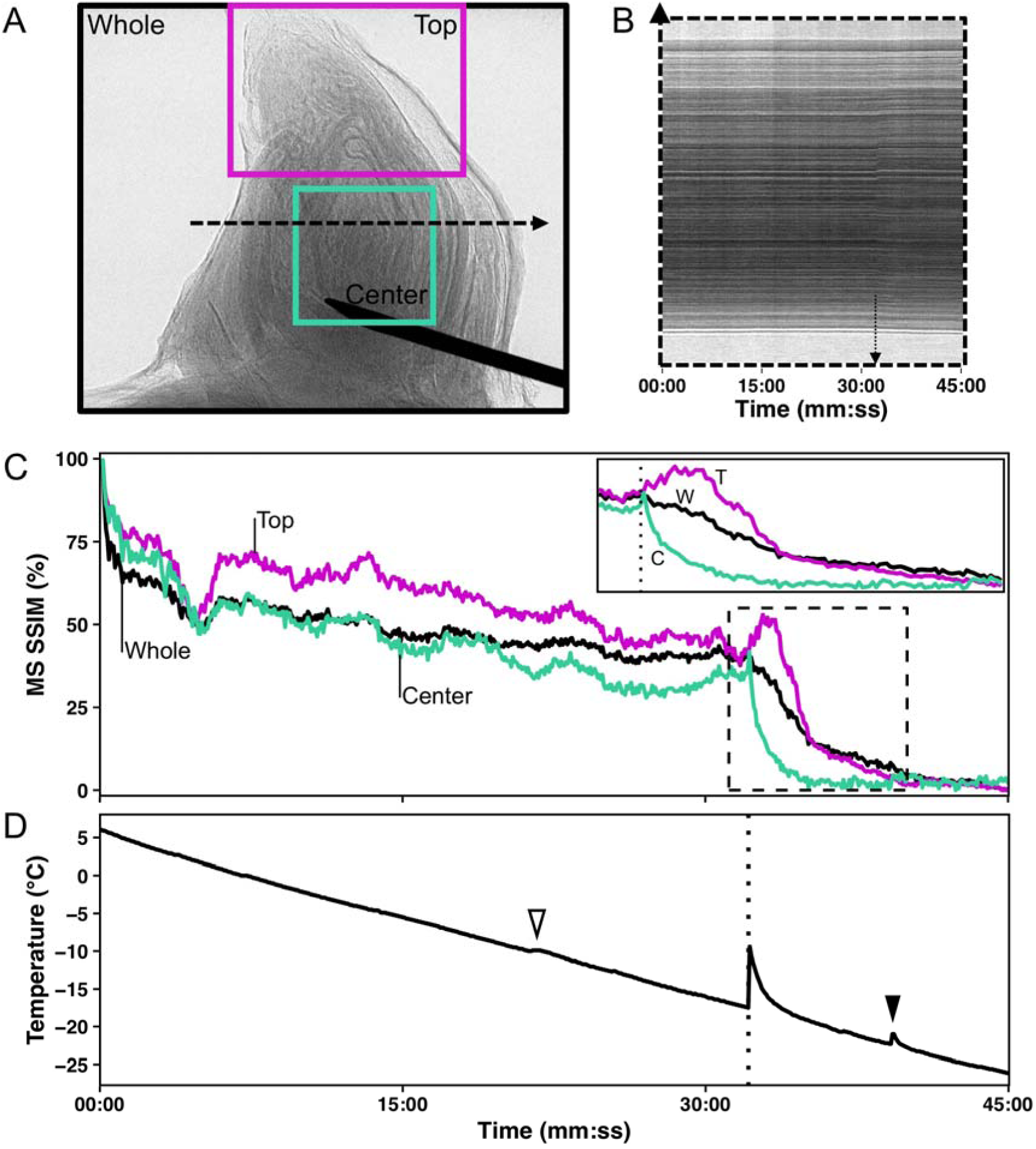
Characteristics of freezing in a bud of *Vitis amurensis* stored at 4 °C (see Supplementary Video S2). (A) Still image of bud at start of freezing; Black (whole image), magenta (top of the bud), and cyan (center of the bud) show areas analyzed; dashed line through center of the bud shows pixels used to build kymograph. (B) Kymograph resulting from line of pixels in the center of the bud; arrows show the start of freezing. (C) Normalized multi-scale structural similarity index (MS SSIM) for three areas in (A); dashed box is shown expanded in the inset, dotted line marks the start of freezing event. (D) Temperature profile measured by thermocouple inside the bud; dotted line marks the start of freezing event, open arrowhead shows slight lag, closed arrowhead shows secondary bud exotherm.

**Video S1.** X-ray phase contrast imaging of *Vitis riparia* bud during freezing. Bud had been exposed to forcing conditions for 11 days (same bud from Figs. 2D and 6). Time stamp on the left in mm:ss format. Expected (blue background, lower) and measured (red background, upper) temperatures in the bud. Expected temperature was calculated based on linear regression of inner bud temperature during cooling while freezing did not occur. Freezing begins at −9.4 °C / 28:41.

**Video S2.** X-ray phase contrast imaging of *Vitis amurensis* bud during freezing. Bud had not experienced forcing conditions (same bud from Figure 4A). Time stamp on the left in mm:ss format. Expected (blue background, lower) and measured (red background, upper) temperatures in the bud. Expected temperature was calculated based on linear regression of inner bud temperature during cooling while freezing did not occur. Freezing begins at −17.5 °C / 22:10.

**Video S3.** X-ray phase contrast imaging of *Vitis vinifera* bud during freezing. Bud had been exposed to forcing conditions for 2 days (same bud from Figure 3B). Time stamp on the left in mm:ss format. Expected (blue background, lower) and measured (red background, upper) temperatures in the bud. Expected temperature was calculated based on linear regression of inner bud temperature during cooling while freezing did not occur. Freezing begins at −18.3 °C / 33:30.

